# Kinase Suppressor of RAS 1 (KSR1) maintains the transformed phenotype of BRAFV600E mutant human melanoma cells

**DOI:** 10.1101/2022.08.16.504086

**Authors:** Zhi Liu, Aleksandar Krstic, Ashish Neve, Nora Rauch, Kieran Wynne, Hilary Cassidy, Amanda McCann, Emma Kavanagh, Brendan McCann, Alfonso Blanco, Jens Rauch, Walter Kolch

**Author notes:** joint corresponding authors **Corresponding authors:** Walter Kolch, Systems Biology Ireland (SBI), University College Dublin, Dublin 4, Ireland., Jens Rauch, Systems Biology Ireland (SBI), University College Dublin, Dublin 4, Ireland.

## Abstract

KSR1 is a scaffolding protein for the RAS-RAF-MEK-ERK pathway, which is one of the most frequently altered pathways in human cancers. Previous results have shown that KSR1 has a critical role in mutant RAS mediated transformation. Here, we examined the role of KSR1 in mutant BRAF transformation. We used CRISPR/Cas9 to knock out KSR1 in a BRAFV600E transformed melanoma cell line. KSR1 loss produced a complex phenotype characterized by impaired proliferation, cell cycle defects, decreased transformation, decreased invasive migration, increased cellular senescence, and increased apoptosis. To decipher this phenotype, we used a combination of proteomic ERK substrate profiling, global protein expression profiling, and biochemical validation assays. The results suggest that KSR1 directs ERK to phosphorylate substrates that have a critical role in ensuring cell survival. The results further indicate that KSR1 loss induces the activation of p38 Mitogen-Activated Protein Kinase (MAPK) and subsequent cell cycle aberrations and senescence. In summary, KSR1 function plays a key role in oncogenic BRAF transformation.

## Introduction

The RAS-RAF-MEK-ERK pathway (hereafter called ERK pathway) is a central signalling pathway of the cell. It is mutationally altered in 30-40% of all human cancers and may be hyperactivated in the majority of cancers, due to crosstalk with other pathways (1). The ERK pathway has a bewildering array of functions (2), and this versatility is tightly coordinated by activation dynamics and scaffolding proteins (3,4). The scaffold protein Kinase Suppressor of RAS 1 (KSR1) has emerged as a major facilitator of normal and oncogenic RAS signalling by binding all three kinases in the pathway, i.e. RAF, MEK, and ERK. Originally, KSR1 was considered a platform that facilitates RAF phosphorylating MEK, and MEK phosphorylating ERK by bringing the kinases in physical proximity. However, a more nuanced view of KSR1 functions is emerging (5). KSR1 not only binds these kinases, but also regulates their activation. For instance, MEK binding to KSR1 stimulates its binding to BRAF resulting in allosteric activation of BRAF’s kinase activity towards MEK (6). Similarly, KSR1 preferentially binds ERK dimers and directs them to cytosolic substrates (7,8).

The perhaps most intriguing finding is that KSR1 knockout mice are healthy, but resistant to oncogenic RAS tumorigenesis (9). While this protection may not be complete in all cancer types (10), it has sparked substantial interest in finding out more about KSR1 functions in oncogenic transformation. As a result, we now know that KSR1 regulates several aspects of oncogenic RAS and RAF transformation, including cell proliferation (11), apoptosis (12), senescence (13,14), and the epithelial-mesenchymal transition (EMT) (15). Most of these KSR1 functions facilitate RAS transformation, and KSR1 has become a plausible drug target for combating RAS driven cancers (16). However, how KSR1 may contribute to transformation by mutant, oncogenic BRAF is not well understood. Therefore, we knocked out KSR1 in a BRAFV600E driven melanoma cells. The knockout resulted in a complex phenotype with features of cell cycle aberration, senescence and enhanced apoptosis. Analysis of the molecular mechanisms suggests a multi-layered mechanism that includes KSR1 controlling ERK substrate specificity.

## Materials and Methods

### Cells

SK-MEL-239 cells (RRID:CVCL_6122) were obtained from Dr Poulikos Poulikakos, Memorial Sloan-Kettering Cancer Center, New York, USA, and cultured in RPMI-1640 complete medium (Gibco, Cat# 21875034) containing 10% FBS and 2mM L-Glutamine (Gibco, Cat# 25030081). Cells were authenticated using the AmpFlSTR® Identifiler® Plus PCR Amplification Kit (ThermoFisher, Cat# A26182; Table S1).

### CRISPR/Cas9 knockout

Three crRNAs (Fig. 1A) that target exon 5 of KSR1 (Ensembl transcript ID ENST00000509603.6) were designed using GeneArt (https://www.thermofisher.com/ie/en/home/life-science/genome-editing/geneart-crispr/geneart-crispr-search-and-design-tool.html) and cloned into the GeneArt™ CRISPR Nuclease Vector with OFP (orange fluorescent protein) Reporter (ThermoFisher, Cat# A21174). Twenty-four hours after transfection, 288 single cells expressing OFP were sorted for each crRNA using a FACSAria III instrument (BD Biosciences). Twenty-four surviving single cell clones for each crRNA were tested for indels using the GeneArt™ Genomic Cleavage Detection Kit (ThermoFisher, Cat# A24372). Positive clones were Sanger sequenced, and the sequence containing mixed base calls from different KSR1 alleles was decomposed by the Synthego webtool (https://ice.synthego.com/#/) (Fig. S1).

**Figure 1.**
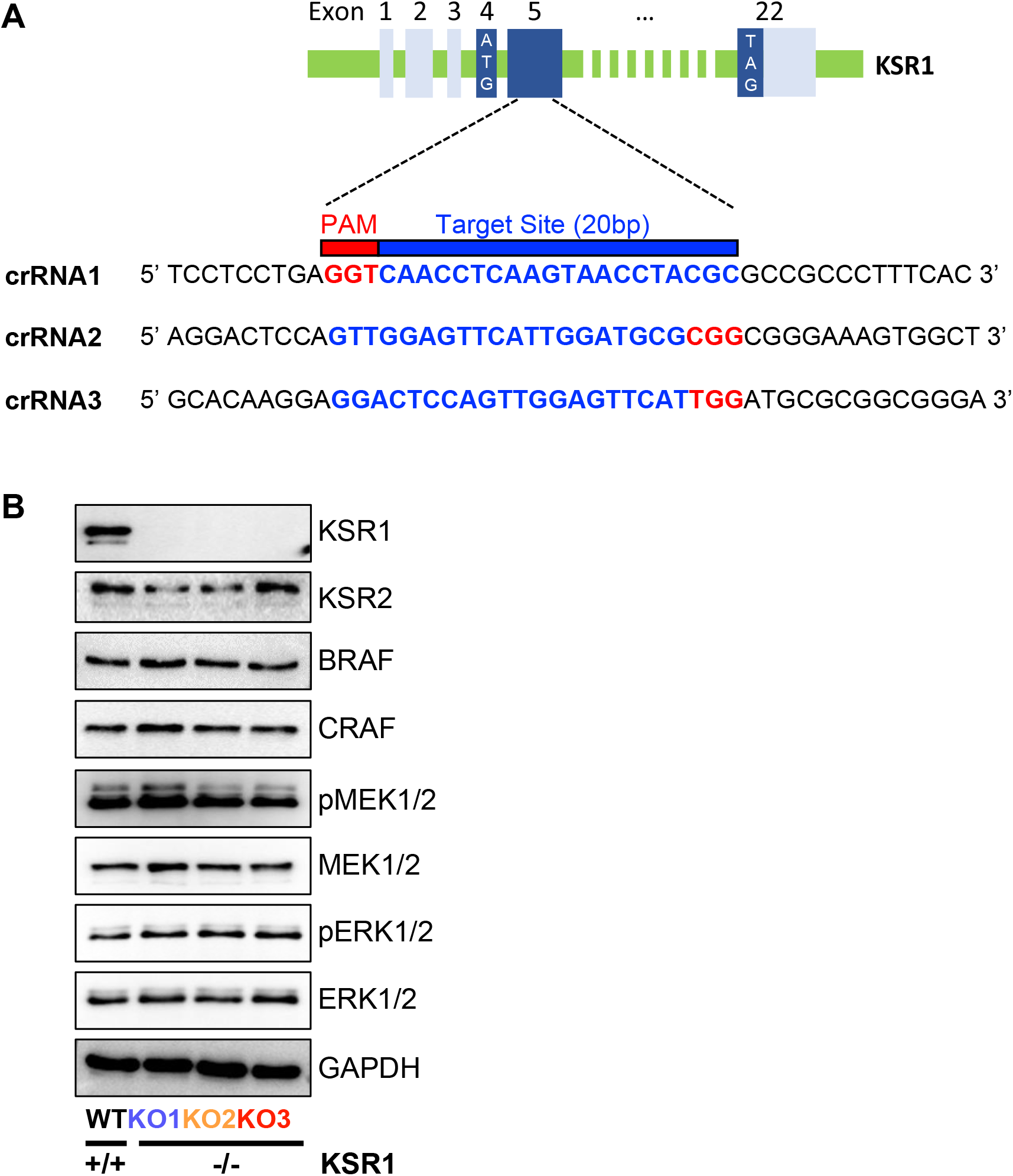
Knockout of KSR1 in BRAFV600E mediated SK-MEL-239 cells does not impact bulk RAF-ERK signalling. **(A)** Schematic of KSR1 exon 1-5 structure and sequence of sgRNAs targeting exon 5. Untranslated exons are shown in light blue, translated exons in dark blue. Translation start (ATG) and stop (TAG) codons are indicated. **(B)** KSR1/2 and RAF-MEK-ERK pathway proteins were detected by Western blotting in wildtype (WT) cells and three KSR1 knock-out clones (KO1-3). MEK and ERK activation was assessed using phosphospecific antibodies (pMEK and pERK).

### Cell proliferation

was measured using the Cell Counting Kit-8 (Sigma-Aldrich, Cat# 96992-500) according to the manufacturer’s instructions.

### Cell senescence

was measured using the Senescence β-Galactosidase Staining Kit (Cell Signaling Technologies, Cat# 9860). Senescent (β-Galactosidase positive) cells were counted manually from three randomly taken fields.

### Anchorage independent growth

was measured using soft agar assays as previously described (17). Colonies were stained with 0.005% crystal violet solution for two hours and manually enumerated.

### Cell cycle analysis

Cells at 80% confluence were collected and resuspended in 1 ml phosphate-buffered-saline (PBS) containing 10μM BrdU (Bio-Sciences, Cat# 556028) for 1 hour. Cells were then fixed in 70% ethanol for 30 minutes and labelled with 10μg/ml propidium-iodide (Sigma, Cat# P4864) for 30 minutes prior to cell cycle analysis using a BD Accuri C6 Flow Cytometer® (BD Biosciences).

### Cell apoptosis

When the cells reached 70 - 80% confluence, they were gently washed with PBS and resuspended in PBS containing 0.1μM YO-PRO-1 (ThermoFisher, Cat# Y3603) and 1μM propidium-iodide (Sigma, Cat# P4864) for 20 minutes prior to analysis on a BD Accuri C6 Flow Cytometer®.

### Trans-well cell migration

was measured using Corning® Transwell® 8μm pore polycarbonate membrane cell culture inserts (Merck, Cat# CLS3422). 1×10^6^ cells in serum-free RPMI medium were added to the inserts with RPMI containing 10% FBS serving as chemoattractant in the bottom chamber. After 24 hours cells migrating through the membrane were fixed with 70% ethanol, stained with Giemsa, and counted.

### Immunocytochemistry

Cells were fixed with ice-cold methanol (Sigma-Aldrich, Cat# 24229) and stained with Ki67 antibody (ThermoFisher, Cat# MA5-15690, RRID:AB_10979995, 1:1000 dilution), using the Novolink Polymer Detection kit (Leica, Cat# RE7140-CE). Cells were counterstained with haematoxylin (Reagecon, Cat# RBA-4201-00A).

### Western blotting

Proteins were separated by SDS-polyacrylamide gel electrophoresis and transferred to a polyvinylidene difluoride membrane using the XCell SureLock® Mini-Cell chamber wet transfer system according to the manufacturer’s instructions (ThermoFisher). Membranes were blocked with 5% non-fat dry milk in TBST (20 mM TrisHCl, pH 7.4; 150 mM NaCl, 0.1% Tween 20) for 30 minutes and washed 3 × 5 minutes in TBST buffer. Then, membranes were incubated overnight with primary antibody in TBST with 4% bovine serum albumin, washed 3 × 5 minutes in TBST buffer, and incubated with secondary antibody (horse radish peroxidase conjugated) in TBST with 5% non-fat dry milk for 1 hour. After three 5-minute washes, the membrane was briefly rinsed with water and developed with Pierce-ECL (enhanced chemiluminescence) reagent (ThermoFisher, Cat# 32109). Bands were visualised using the ChemiImager (Advanced Molecular Vision accompanied with Chemostar software) or iBright™ CL750 Imaging System (Invitrogen). To re-use membranes, antibodies were removed by incubation in stripping buffer for 15 minutes (0.2 M glycine, pH 2.5, 1% SDS).

### Antibodies used for Western blotting

were from the following vendors: Cell Signaling Technology, KSR1 (Cat# 4640, RRID:AB_10544539), pMEK1/2 (Cat# 9121, RRID:AB_331648), MEK1/2 (Cat# 9122, RRID:AB_823567), GAPDH (Cat# 2118, RRID:AB_561053), HSP90 (Cat# 4877, RRID:AB_2233307), p-p38 (Cat# 9211, RRID:AB_331641), p38 (Cat# 9212, RRID:AB_330713), p53 and p53 phospho-forms pS15 and pS392 (Phospho-p53 Antibody Sampler Kit #9919, RRID:AB_330019), Rb1 pS780 (Cat# 9307, RRID:AB_330015), E-Cadherin (Cat# 3195, RRID:AB_2291471), MITF (Cat# 97800S, RRID:AB_2800289), p14ARF (Cat# 74560S, RRID:AB_2923025), PDCD4 (Cat# 9535, RRID:AB_2162318), horse radish peroxidase-linked anti-mouse IgG (Cat#7076, RRID:AB_330924), horse radish peroxidase-linked anti-rabbit IgG (Cat#7074, RRID:AB_2099233); Santa Cruz, KSR2 (Cat# sc-100421, RRID:AB_1124518), BRAF (Cat# sc-5284, RRID:AB_626760), CRAF (Cat# sc-133, RRID:AB_632305); p21 (Cat# Sc-6246, RRID:AB_628073), CDK4 (Cat# sc-23896, RRID:AB_627239), CDK6 (Cat# sc-7961, RRID:AB_627242), Caveolin 1 (Cat# sc-894, RRID:AB_2072042); Sigma-Aldrich, pERK1/2 (Cat# M8159, RRID:AB_477245), ERK1/2 (Cat# M5670, RRID:AB_477245); BD Biosciences, p16INK4A (Cat# 550834, RRID:AB_2078446); Abcam, TYRP1 (Cat# Ab235447, RRID:AB_2923026).

### Immunoprecipitation

Cells were lysed in 20mM Tris-HCl (pH 7.5), 150mM NaCl, 0.5% NP-40, 1mM EDTA and 1mM EGTA containing protease inhibitor cocktail (cOmplete™ Mini Protease Inhibitor Cocktail, Roche Diagnostics, Cat# 11836170001), and phosphatase inhibitor cocktail (PhosSTOP, Roche Diagnostics, Cat# 4906837001). Cell lysates were cleared by centrifugation at 15,000xg at 4°C for 10 minutes. The protein concentration of the lysates was determined by Pierce® BCA Protein Assay (ThermoFisher, Cat# 23225). Potential ERK substrates were immunoprecipitated using Phospho-MAPK/CDK Substrates sepharose beads (Cell Signaling, Cat# 5501) from cell lysates at 4°C for 6 hours. The immunoprecipitates were washed 3x with lysis buffer and processed for mass-spectrometry as described previously (18).

### Mass Spectrometry analysis

of immunoprecipitates was performed as previously reported (Turriziani et al., 2014). Total protein expression profiling used the SP3 method (19). The raw data were analysed by MaxQuant and Perseus (20).

### Lysate based proteomics

Cells were resuspended in 100µl 8M urea, 50mM Tris-HCl pH 8.0, supplemented with cOmplete™ Mini Protease Inhibitor Cocktail (Roche Diagnostics, Cat# 11836170001), and phosphatase inhibitor cocktail (PhosSTOP, Roche Diagnostics, Cat# 4906837001). Samples were sonicated for 2 × 9 seconds at a power setting of 15% to disrupt cell pellets (Syclon Ultrasonic Homogenizer). Sample were reduced by adding 8mM dithiothreitol (DTT) for 60 minutes and subsequently carboxylated using 20mM iodoacetamide for 30 minutes in the dark while mixing (Thermomixer 1,200rpm, 30°C). The solution was diluted with 50mM Tris-HCl pH 8.0 to a final urea concentration of 2M. Sequencing Grade Modified Trypsin (Promega Cat# V5111) was resuspended in 50mM Tris-HCL at a concentration of 0.5ug/µl and added at a 1:100 enzyme to protein ratio. Samples were digested overnight with gentle shaking (thermomixer 850rpm, 37°C). The tryptic digest was terminated by adding formic acid to 1% final concentration and samples were desalted using C18 HyperSep™ SpinTips (ThermoFisher, Cat# 60109-412).

### Geneset Enrichment analysis

(GSEA) was performed using EnrichR (21).

#### Statistics

Two-tailed, paired or unpaired Student’s T-Test was performed to analyse the significance of difference between two groups. GraphPad Prism version 5.01 (RRID:SCR_002798) was used to create graphs. Error bars represent standard-error-of the-mean (SEM); 1-3 asterisks indicate significance at the 0.05, 0.01, or 0.001 probability level, respectively; *n*.*s*. indicates non-significant. As the work focuses on the molecular mechanistic analysis of KSR1 loss in cell lines, blinding, power analysis, randomisation, and considerations regarding differences between males and females were not required for the study.

#### Data Availability Statement

The data supporting the findings of this study are currently submitted to PRIDE.

## Results

### Knocking out KSR1 in BRAFV600E mutated SK-MEL-239 cells does not impact bulk RAF-ERK signalling

To knock out KSR1 gene expression in SK-MEL-239 cells, we used the CRISPR/Cas9-OFP system with three crRNAs (22) that target exon 5 of KSR1 (Figs. 1A, S1A). This exon is common to different KSR1 splice variants and located close to the start of the coding sequence. Its disruption is expected to result in a complete loss of KSR1 protein expression. After isolating successfully transfected, i.e. OFP expressing, cells, KSR1 knockout clones were identified by Genomic Cleavage Detection (GCD) assays and Sanger sequencing (Fig. S1B). We selected three clones with homozygous indels in KSR1 exon 5 that cause a complete loss of KSR1 protein expression as detectable by Western blotting (Fig. 1B). There was no compensatory upregulation of KSR2 expression, and the KSR1 knockout did not affect the protein levels of BRAF, CRAF, MEK or ERK. Interestingly, there was also no effect on the activation of MEK and ERK, suggesting that KSR1 function is not required to sustain MEK-ERK activity in these cells. To ensure the lineage fidelity, we genotyped the parental and KSR1 knockout cells finding that they all retain the same genotype (Table. S1).

### The biological phenotype of KSR1 loss

In order to test the biological consequences of the KSR1 knockout, we assayed different biological traits. KSR1 knockout cells proliferated significantly slower than the parental cells (Fig. 2A). Cell cycle analysis showed that KSR1 loss did not prevent cells from exiting interphase (G0/G1) but retarded their progression through late S and G2/M phase (Figs. 2B, S2) suggesting that KSR1 function is needed to complete the cell cycle after DNA replication.

**Figure 2.**
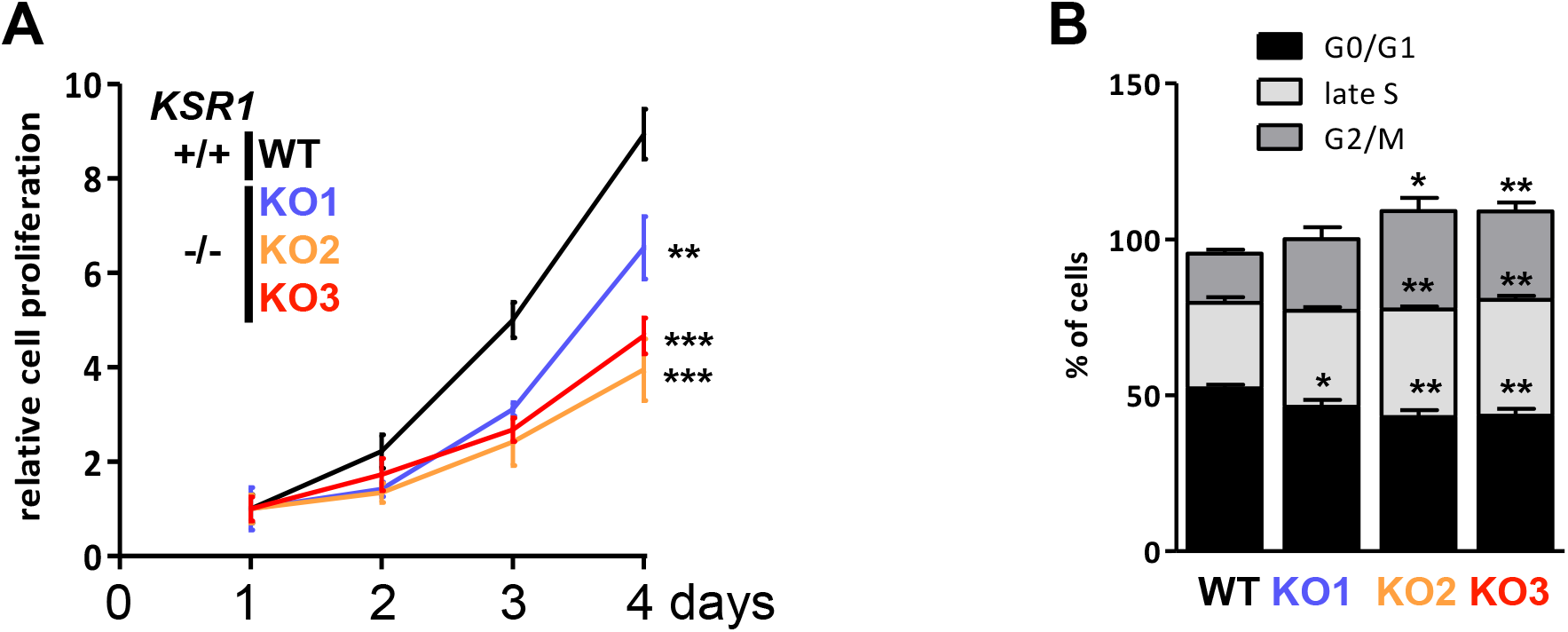
KSR1 loss decreases proliferation by retarding cell cycle progression through late S and G2/M phase. **(A)** Cell proliferation. **(B)** Cell cycle analysis.

We noticed that KSR1^-/-^ cultures contain large, flat cells that resemble the phenotype of senescent cells. Performing a stain for acidic β-galactosidase confirmed an increase in the number of senescent cells in KSR1 KO1-3 clones (Fig. 3A). The expression of the proliferation marker Ki67 was attenuated in the phenotypically senescent cells (Fig. 3B). Interestingly, most of the non-proliferative, acidic β-galactosidase positive cells were multinucleated, further supporting the interpretation of the cell cycle data that KSR1^-/-^ cells can replicate DNA but are unable to complete mitosis and cell fission. Cells that arrest in mitosis for a prolonged time typically die by apoptosis or exit mitosis without dividing, causing a multinucleated phenotype (23). Indeed, all KSR1 KO clones showed increased rates of apoptosis (Fig. 3C) suggesting that the decrease in cell proliferation is caused by a combined increase in senescence and apoptosis.

**Figure 3.**
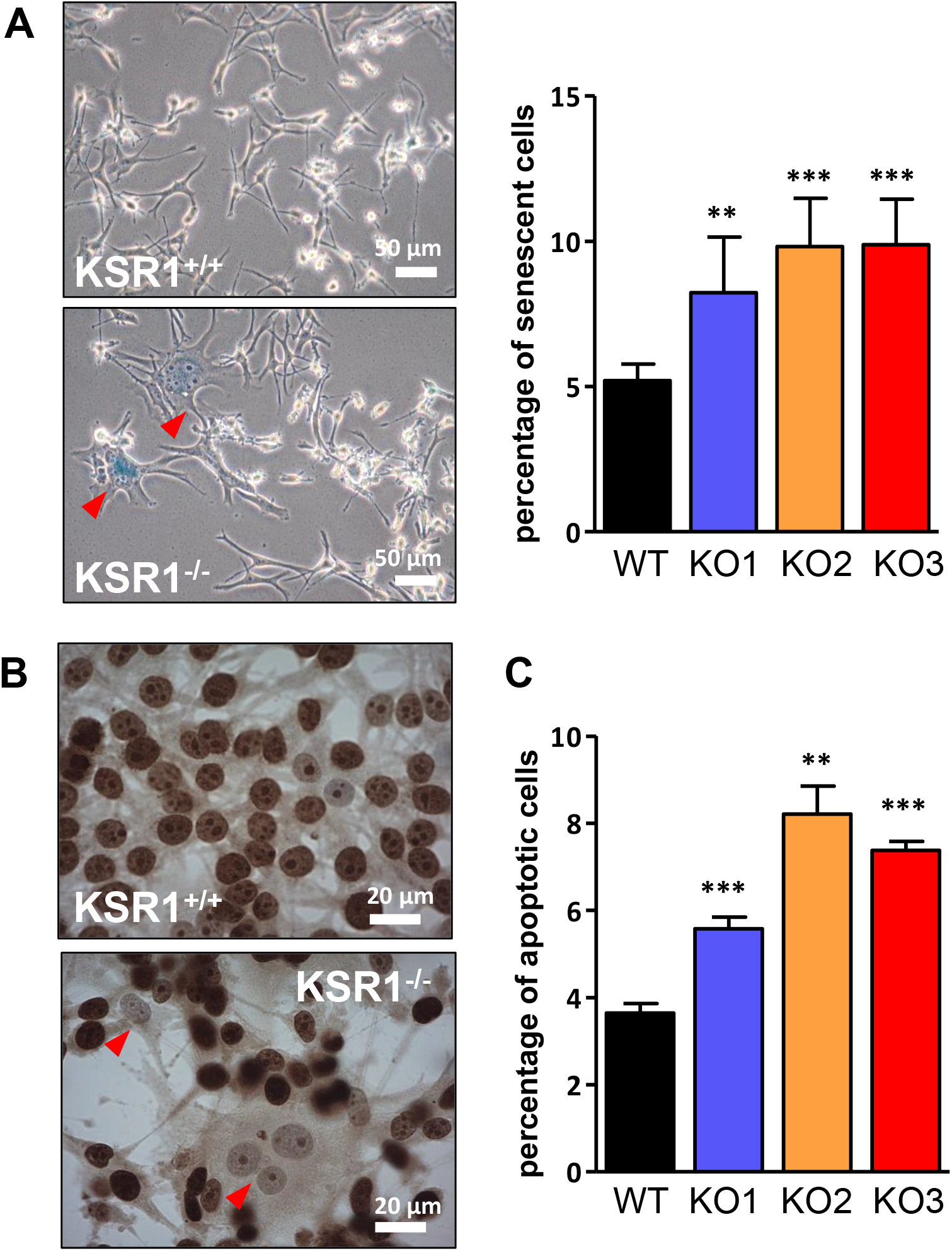
KSR1 loss increases senescence and apoptosis. **(A)** Increased expression of the senescence marker acidic β-galactosidase in KSR1^-/-^ cells. Left panel, representative acidic β-galactosidase stains; right panel, quantification of β-galactosidase activity; red arrowheads indicate β-galactosidase positive cells. **(B)** Ki-67 stain indicating proliferative cells. Red arrowheads indicate multinucleated cells with senescent morphology. **(C)** Percentage of apoptotic cells measured by the YO-PRO™-1 Iodide assay.

These results indicated that KSR1 may have a role in sustaining the transformed phenotype of melanoma cells. Therefore, we tested the effects of KSR1 knockout on the ability of cells to grow in 3D soft agar cultures, which is a reliable *in vitro* indicator of tumorigenicity *in vivo* (24). KSR1^-/-^ cells failed to grow in soft agar, whereas parental cells formed readily visible colonies (Fig. 4A). Similarly, KSR1 knockout severely compromised the ability of SK-MEL-239 cells to invasively migrate through a Transwell membrane (Fig. 4B).

**Figure 4.**
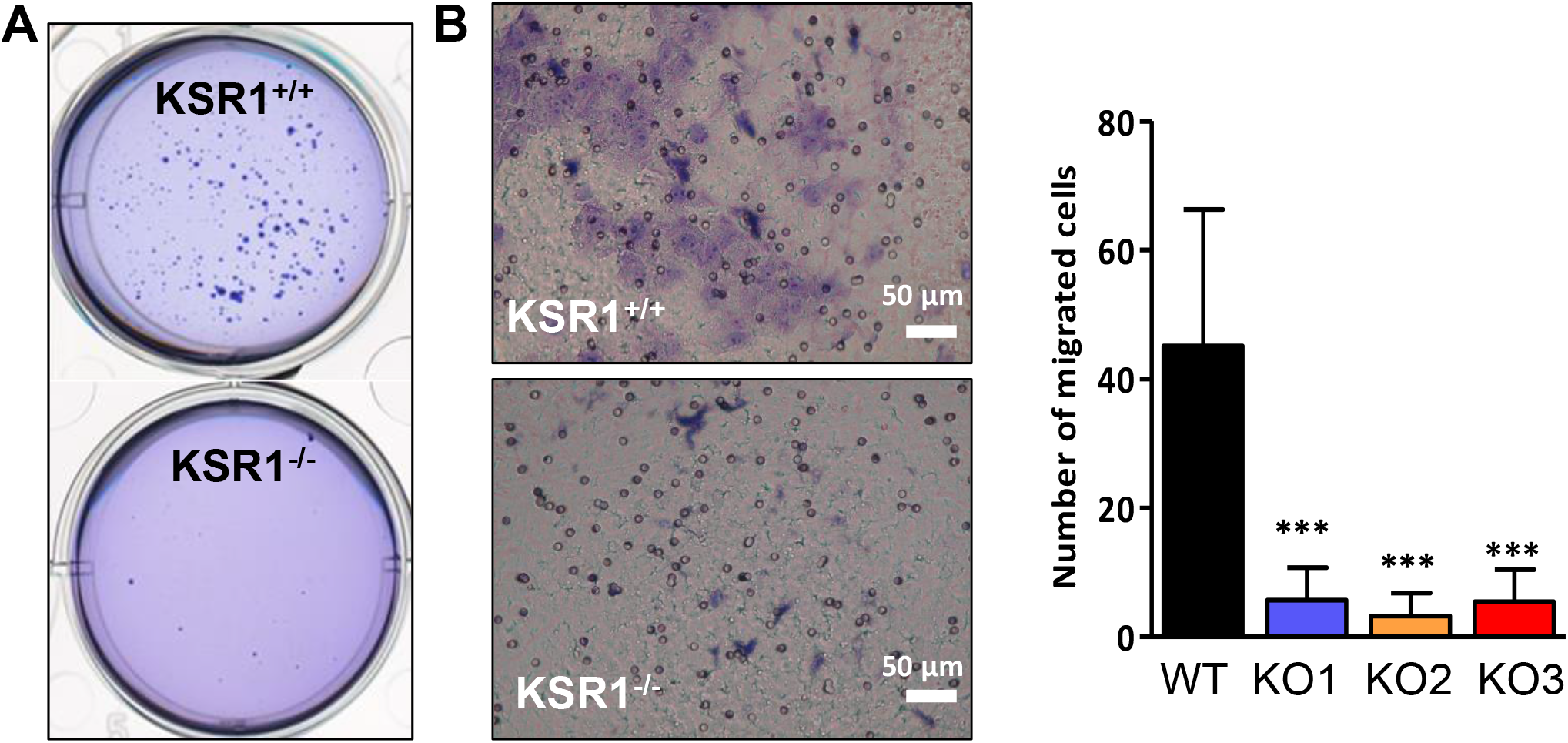
KSR1 loss inhibits growth in soft agar and invasive migration. **(A)** A representative assay showing that KSR1^-/-^ cells fail to form colonies in soft agar. **(B)** Transwell migration assay. Cells were stained with Giemsa, and cells able to migrate through the membrane were counted. Left panel, representative image; right panel, quantification of cells that have migrated through the membrane.

In conclusion, KSR1 loss interferes not only with cell proliferation and cell cycle progression, but also with several traits of oncogenic transformation including the ability of anchorage independent growth in soft agar and invasive migration.

### ERK-substrateomics

The above results suggested that KSR1 plays an important role in maintaining the transformed state of BRAF mutant melanoma cells. As ERK activation is considered a main effector of mutant BRAF signalling, we re-examined the role of ERK in more depth. Given the lack of impact of KSR1 loss on global ERK activity (Fig. 1B), we hypothesized that KSR1 may direct ERK to specific substrates rather than being required for general ERK activation. ERK fulfils its pleiotropic biological functions via almost 500 bona fide substrates (25), whose phosphorylation plausibly needs to be selective in order to achieve specific biological outcomes. Therefore, we assessed the impact of KSR1 loss on the phosphorylation of ERK substrates.

For this, we enriched ERK substrates using an antibody that recognizes sites phosphorylated by ERK (P-X-pS-P and pS-X-R/K), and identified and quantified the immunoprecipitated ERK substrates by mass spectrometry (MS) (Fig. 5A). Consistent with the observation that KSR1 knockout did not impact global MEK and ERK activation, the pattern of ERK phosphorylated proteins resolved by gel electrophoresis was highly similar between WT and KSR1 KO cells (Fig. 5B). However, MS analysis revealed a small number of ERK substrates that were differentially phosphorylated (Figs. 5C, D; Table S2). Of 399 proteins specially immunoprecipitated (i.e. enriched >2 fold over a control immunoprecipitation with an isotype matched IgG) in KSR1 KO1-3 cells, 85 were known ERK substrates (26). Analysing differences between parental and KSR1^-/-^ cells using a fold change of >2 and p-value of <0.05 as cut-off for differential phosphorylation, we identified 29-34 substrates showing enhanced and 28-33 substrates showing decreased phosphorylation in the KSR1 KO1-3 clones versus parental cells. This finding supports our hypothesis that KSR1 may direct ERK to specific substrates. Interestingly, only 6 up- and 4 downregulated substrate phosphorylations were shared between all 3 KSR1 knockout clones suggesting that cells can adapt to KSR1 loss via different mechanisms that share common core processes (Fig. 5C, D).

**Figure 5.**
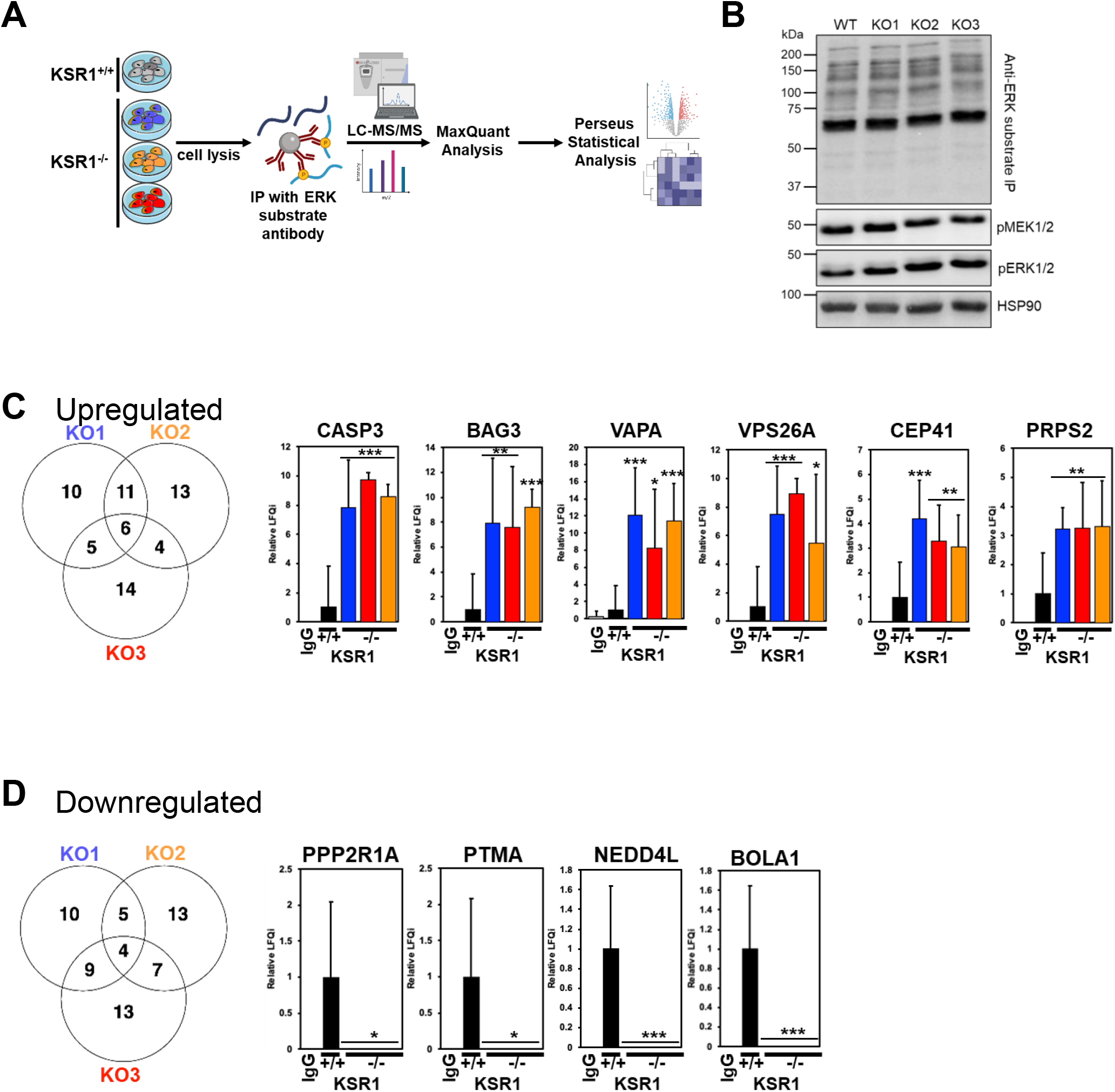
ERK substrateomics. **(A)** Workflow. **(B)** Western blot of ERK substrate immunoprecipitation (IP) stained with ERK substrate antibody. Lysates were stained for activated MEK (pMEK) and ERK (pERK). HSP90 served as loading control. **(C**,**D)** Proteins whose phosphorylation was up- or downregulated in KSR^-/-^ cells. The changes shared by KSR KO1-3 clones are shown as bar graphs from 3 biological replicates.

These core adaptations include a very significant increase in the phosphorylation of caspase 3, BAG3 (Bcl-2-associated athanogene 3), VAPA (VAMP Associated Protein A), VPS26A (Vacuolar Protein Sorting-Associated Protein 26A), CEP41 (Centrosomal Protein 41), and PRPS2 (Phosphoribosyl Pyrophosphate Synthetase 2). Caspase 3 integrates both extrinsic and intrinsic apoptosis pathways and is a key effector of apoptosis (27). Caspase 3 has an ERK docking site, and ERK can activate caspase 3 (28), which could help explaining the enhanced apoptosis in KSR1^-/-^ cells. BAG3 is a multifunctional protein that is involved in protein folding, autophagy and apoptosis (29). Interestingly, ERK phosphorylation neutralizes its protective function against oxidative stress induced apoptosis (30) suggesting that the enhanced BAG3 phosphorylation common to KSR^-/-^ cells can contribute to their increased apoptosis rates. VAPA and VPS26A function in vesicle transport (31,32). CEP41 is a centrosomal protein that regulates the function of cilia (33). PRPS2 functions in the deoxynucleotide synthesis pathway and its overexpression stimulates the proliferation and metastatic capacity of melanoma cells (34). Proteins whose phosphorylation at ERK target sites was significantly downregulated in all KSR1^-/-^ clones include PPP2R1A (Protein Phosphatase 2 Regulatory Subunit A α), PTMA (Prothymosin α), NEDD4L (NEDD4 Like E3 Ubiquitin Protein Ligase), and BOLA1 (BolA Family Member 1). PPP2R1A is a subunit of the serine/threonine phosphatase PP2, which directs PP2 to specific substrates, and functions as tumour suppressor in endometrial cancer (35). PTMA is an immunomodulatory protein that can enhance T cell responses to tumours (36). On the other hand, PTMA expression in melanoma cells enhances their growth and aggressiveness in a preclinical mouse model (37). These different actions could conceivably be dependent on posttranslational modifications, such as phosphorylation. NEDD4L is an E3 ubiquitin ligase, which can be overexpressed in melanoma and promote tumour growth (38). ERK can phosphorylate NEDD4L on S448, and this phosphorylation is reduced in melanoma cells that are resistant to the RAF inhibitor PLX4720 (39). Phosphorylation of this site disrupts substrate binding and is an effective inhibitor of NEDD4L function (40). BOLA1 helps maintaining the mitochondrial redox balance by counteracting the effects of glutathione depletion (41).

Of these 10 proteins only two are listed in the “Compendium of ERK targets” (26), specifically, BAG 3 and NEDD4L. In both cases ERK phosphorylation inhibits their function. The effects are consistent with the phosphorylation changes observed in KSR^-/-^ cells, i.e., the increase of BAG3 phosphorylation reducing cell survival and the decrease in NEDD4L phosphorylation blocking invasive cell migration. The distinct phosphorylation changes in ERK substrates in KSR1 knockout cells require further investigation in dedicated functional studies. They, however, support our hypothesis that KSR1 can direct ERK substrate phosphorylation.

### KSR1 dependent global changes in protein expression

The specific changes in ERK substrate phosphorylation caused by KSR1 knockout prompted us to investigate whether KSR1 knockout also alters protein expression. We used label free quantitative proteomics to profile global protein expression in parental SK-MEL-239 cells and the three KSR1^-/-^ clones (Fig. 6A). The expression of several proteins was differentially regulated between parental and KSR1^-/-^ SK-MEL-239 cells (Fig. 6B, Fig. S3, Table S3). The expression of 36/21 proteins was up/downregulated, respectively. The proteins downregulated in KSR1^-/-^ cells are involved in tetrahydrofolate and pyrimidine (deoxythymidine) synthesis (Fig. 6C), which may contribute to the S-phase delay in KSR1-/- cells (Figs 2B,S2). The upregulated proteins mapped onto signalling pathways for apoptosis, senescence, autophagy and the p53 network among the top hits (Fig. 6C). These mappings correspond well to the observed phenotype of the KSR1^-/-^ cells. While the ERK substrateomics provided plausible explanations for the apoptosis and migration phenotype, the senescence and cell cycle phenotypes remained elusive.

**Figure 6.**
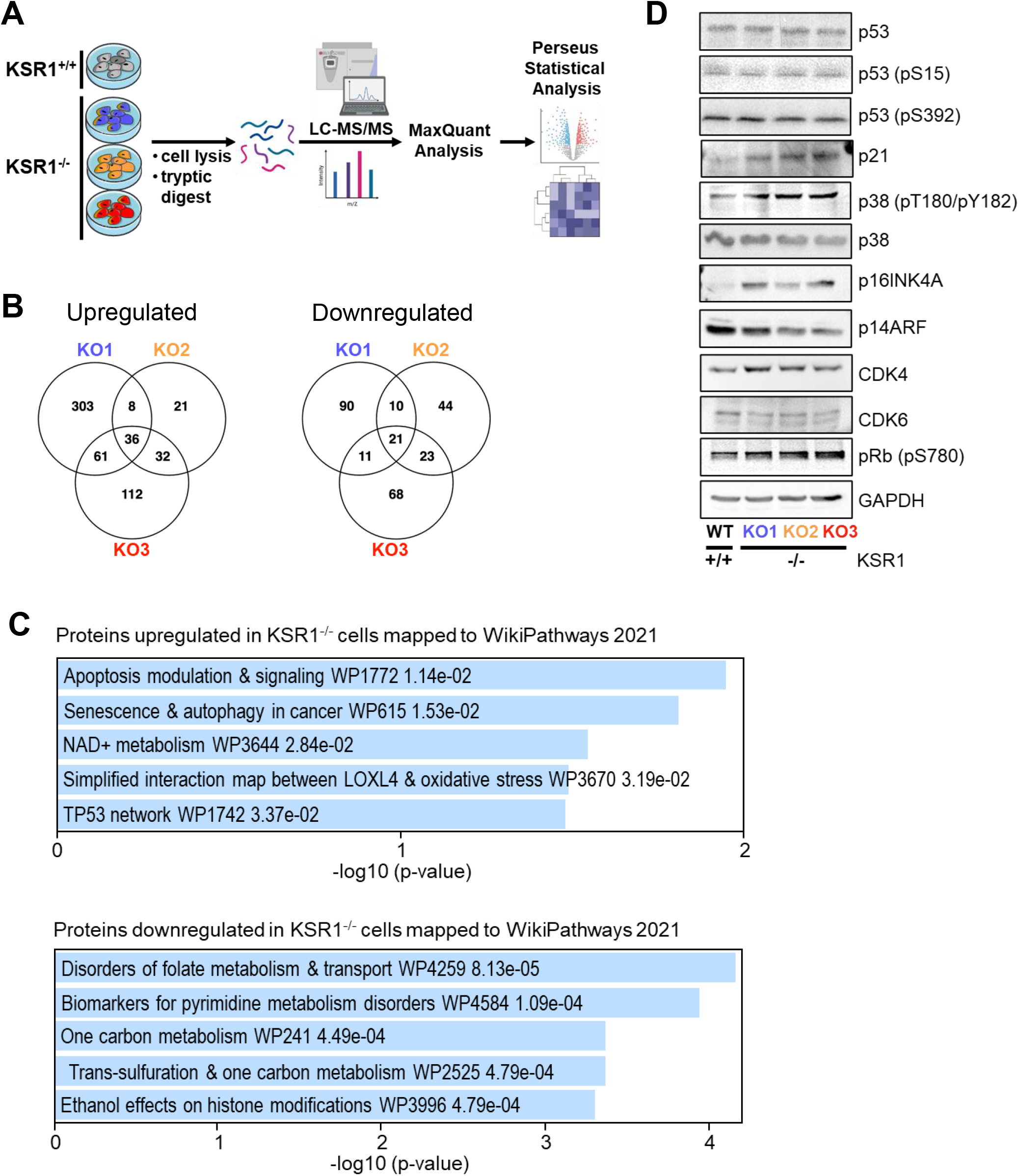
Global proteomic expression profiling. **(A)** Workflow. **(B)** Venn diagram of proteins differentially regulated in the KSR KO1-3 clones vs parental SK-MEL-239 cells. **(C)** ENRICHR analysis of the differentially expressed proteins. The combined score is the log p-value multiplied by the Z-score of the deviation from the expected rank. **(D)** Western blot validation of key protein expression changes found by MS based proteomic expression profiling.

Given that the global expression proteomics highlighted senescence and p53, and that p53 is a major player in both senescence and cell cycle regulation (42), we examined the status of the p53 pathway in the KSR1^-/-^ cells in more detail using Western blot analysis of key proteins (Fig. 6D). These proteins were chosen based on existing knowledge of pathways that connect cell cycle, senescence and p53. Surprisingly, Western blot analysis showed no changes in p53 abundance or phosphorylation of sites that regulate p53 activity. However, the protein expression of the cell cycle inhibitor p21, a classic transcriptional p53 target gene, was upregulated. The p38 kinase can stabilize the p21 mRNA and thereby enhance p21 protein expression independently of p53 (43), and p38 is also implicated in senescence (44). Indeed, p38 was activated in the KSR1^-/-^ cells. The many roles of p38 in senescence induction include activation of p16INK induction (45). The p16INK protein is encoded by the CDKN2A gene, which also encodes the p14ARF tumour suppressor protein. MS analysis showed that p16INK was upregulated in KSR1^-/-^ cells, and this result was confirmed by the Western blot analysis. The p14ARF protein, which regulates p53 protein stability was downregulated in the KSR1^-/-^ cells. This is consistent with the observation that p53 levels do not change in the KSR1^-/-^ cells. The p16INK protein binds to and inhibits the cell cycle kinases CDK4 and CDK6, which promote cell cycle entry by phosphorylating and inactivating the retinoblastoma protein RB1. The expression levels of CDK4 and CDK6 are similar in parental and KSR1^-/-^ cells, suggesting that the KSR1 knockout affects their regulation rather than their expression. Interestingly, phosphorylation of the RB1 protein at S780 was enhanced in the KSR1^-/-^ cells. This phosphorylation is critical for the inactivation of RB1 and progression of cells into S-phase (46). While the enhanced inactivation of RB1 in KSR1^-/-^ cells seems counterintuitive, it fits the observed phenotype. KSR1^-/-^ cells can still synthesize DNA and enter S phase, before being slowed down in late S and G2 phases (Figs. 2B, 3B). S780 can be phosphorylated by CDK4/6 and several other kinases including p38 in the context of proapoptotic signalling (47). Alternatively, at low concentrations p21 serves as a scaffold that promotes the assembly of CDK4/6 complexes with cyclin D enhancing CDK4/6 activity before it inhibits it at high p21 concentrations (48). These possibilities are not mutually exclusive and will be interesting to dissect in future studies.

In addition to these effects on the cell cycle, apoptosis and senescence, we also found protein expression changes that suggest a role for KSR1 in cell differentiation and adhesion (Fig. S3). The expression of the tumour suppressor protein PDCD4 (Programmed Cell Death 4) correlates with good prognosis in melanoma (49), and is upregulated in KSR1^-/-^ cells. Likewise, CAV1 (Caveolin) is slightly overexpressed in KSR1^-/-^ cells. It functions as tumour suppressor in melanoma and restricts cell growth and motility (50). In mouse embryo fibroblasts CAV1 associates with KSR1 and enhances KSR1 functions (14). Similarly, MITF (Melanocyte Inducing Transcription Factor) is upregulated in KSR1^-/-^ cells. MITF is a transcription factor that initiates and maintains the melanocyte lineage (51). By contrast, TYRP1 (Tyrosinase Related Protein 1) protein expression is severely downregulated in KSR1^-/-^ cells. TYRP1 functions in melanin synthesis, although high expression is associated with poor prognosis due to sequestration of the tumour suppressor miRNA-16 (52). Likewise, β-catenin expression is strongly suppressed in KSR1^-/-^ cells. β-catenin is part of the classic WNT signalling pathway and increases tumorigenicity, metastasis and drug resistance in melanoma (53). Interestingly, enhanced WNT signalling in melanoma cells also inhibits T-cell infiltration and response to immunotherapies (54). These molecular changes are largely consistent with the observed phenotypical changes in response to KSR1 knockout. However, further investigations are required to determine the exact roles of these multiple changes in the KSR^-/-^ phenotype.

## Discussion

Our study confirms the emerging intricacy of KSR1 functions (5). Knocking out KSR1 in the BRAFV600E driven melanoma cell line SK-MEL-239 resulted in a complex phenotype that shows features of aberrant cell cycle regulation, enhanced senescence and increased apoptosis. Interestingly, KSR1 seems to support BRAFV600E driven transformation through different functions, which are not fully explainable by known mechanisms. The decrease in proliferation caused by KSR1 knockout is mainly due to a slowing down of S phase exit, and G2 phase completion. Examining the activity of cell cycle checkpoints showed an upregulation of p21 and p16INK4A in KSR1^-/-^ cells. These proteins are classic inhibitors of S phase entry and should decrease the phosphorylation and inactivation of RB1, which controls the G1/S progression. Our results show that RB1 phosphorylation increases in the KSR1^-/-^ cells. This is consistent with the cell being able to replicate DNA but does not explain why they have difficulties progressing through late S and G2 phases. We did not find any changes in the expression of mitotic CDK inhibitors, such as p27, but a recent report suggests that p21 also can control later stages of the cell cycle (55). An alternative and non-mutually exclusive explanation could be that the p38 MAPK, which is activated in KSR1^-/-^ cells, can phosphorylate and inactivate RB1 independently of CDKs (47). Moreover, the decrease in expression of proteins involved in pyrimidine synthesis (Fig. 6C) may decelerate late S phase by causing cells running out of nucleotides for DNA synthesis.

The increase in senescence caused by KSR1 knockout is also unorthodox. KSR1^-/-^ cells show a clear increase in cells with the classic senescent morphology and expression of the classic senescence marker acidic β-galactosidase as well as an increase in the expression of p21 and p16INK. However, they did not show other hallmark features of senescence, such as upregulation of p53, p27 and proteins typical for the senescence-associated secretory phenotype (SASP). As DNA replication occurred in KSR1^-/-^ cells with multinucleated cells appearing that mainly also had senescent appearance, it is possible that senescence is triggered by endoreplication (56). Nevertheless, it is an unorthodox senescence phenotype, as judged by the usual criteria (57).

The clearest explanation can be provided for the increase in apoptosis. KSR1^-/-^ cells exhibit an increase in inactivating BAG3 and presumably activating Caspase 3 phosphorylation. This would remove a protective mechanism and activate an apoptosis executioner molecule, which plausibly could account for the increase in apoptosis in KSR1^-/-^ cells.

How does this all fit together? Within the limitations that more detailed studies of each aspect will be required to fully disentangle the molecular mechanisms underpinning the KSR1 knockout phenotype, we propose the following model (Fig. 7). Our results suggest that KSR1 controls ERK substrate choice. When KSR1 is lost, ERK can activate the executioner Caspase 3 and inactivate the apoptosis antagonist BAG3 to promote apoptosis. In the p53 and RB1 networks the increase in p21 and p16INK could be due to direct effects of the p38 MAPK, which can increase the expression of both proteins (45,47). Thus, our results indicate that KSR1 has a multi-layered role in facilitating transformation by oncogenic BRAF mutants and that some of these traits my lend themselves to therapeutic interference.

**Figure 7.**
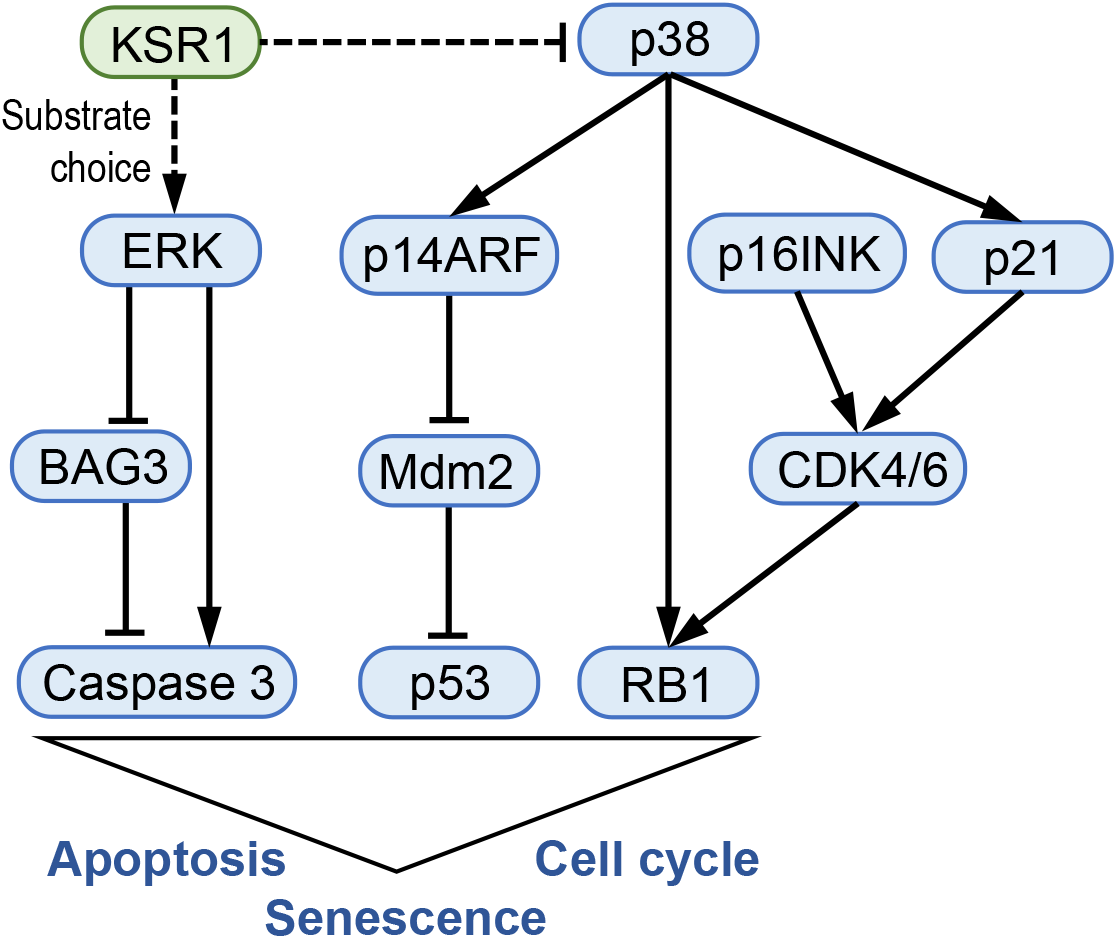
Summary model of KSR1 functions in BRAF mutant melanoma cells. See text for details. Arrows indicate activation; blunt lines indicate inhibition; broken lines indicate that the regulatory mechanism is not entirely clear.

## Acknowledgements

We thank the UCD Conway Core Technologies and the UCD/SBI Comprehensive Molecular Analysis Platform for expert assistance with proteomics and flow cytometry, and Philip Cotter for help with data upload to PRIDE.

## Supplementary Figure Legends

**Supplementary Figure S1.**
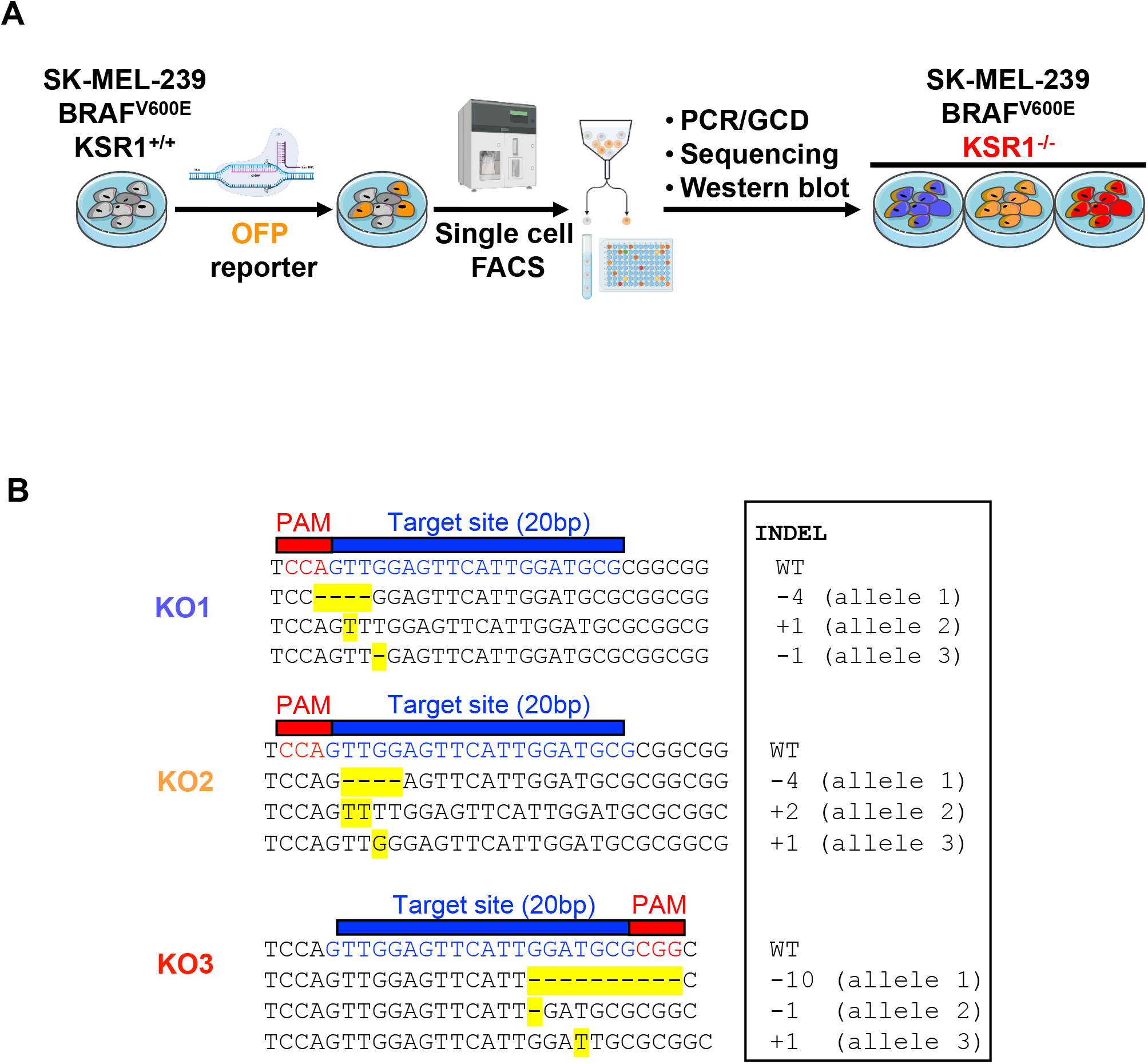
CRISPR/Cas9 mediated knockout of KSR1 in SK-MEL-239 cells. **(A)** Workflow of obtaining KSR1^-/-^ clones. OFP, orange fluorescent protein; FACS, fluorescence activated cell sorting; PCR, polynucleotide chain reaction; GCD, genomic cleavage detection. **(B)** Indels detected in the KSR1 knockout clones KO1-3. Note that SK-MEL-239 cells contain 3 KSR1 alleles. All Indels result in reading frameshifts and truncation of the coding sequence.

**Supplementary Figure S2.**
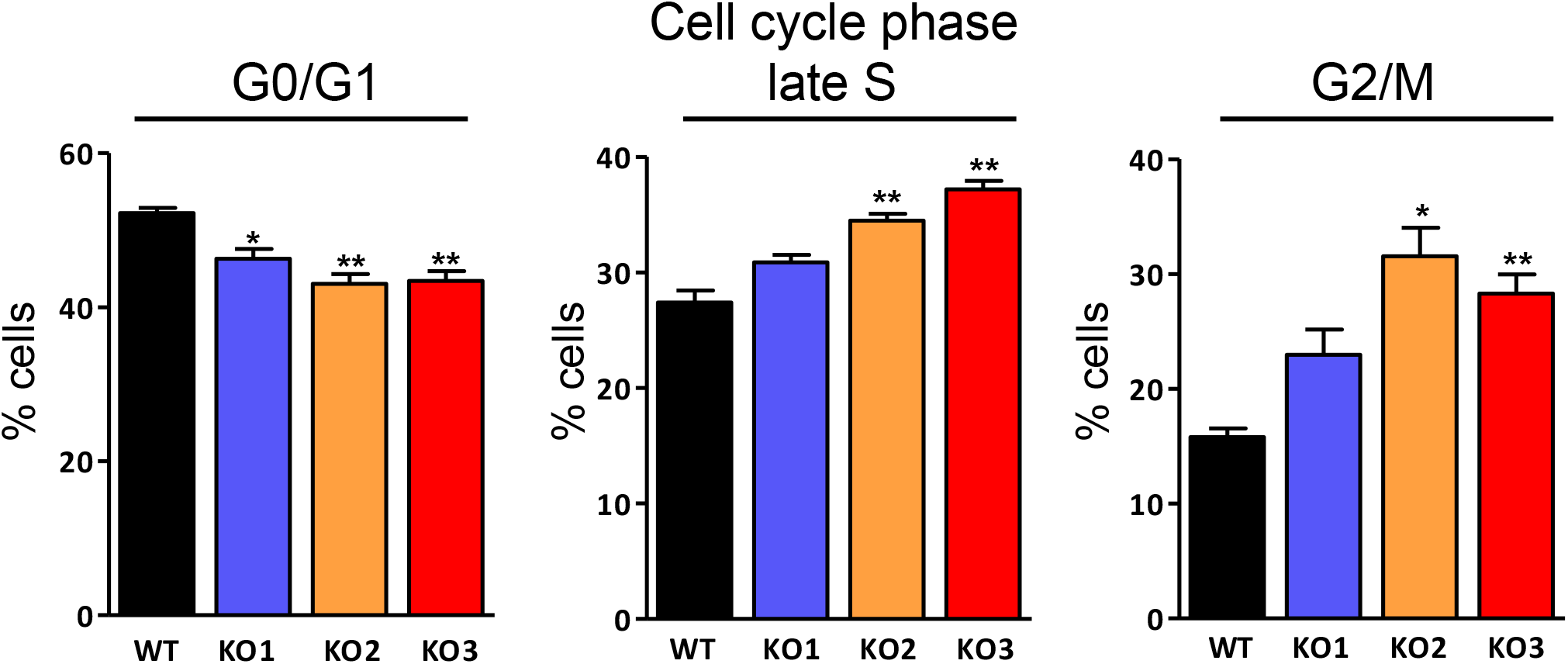
Cell cycle analysis. The data show the individual components of the composite cell cycle analysis presented in Fig. 2B.

**Supplementary Figure S3.**
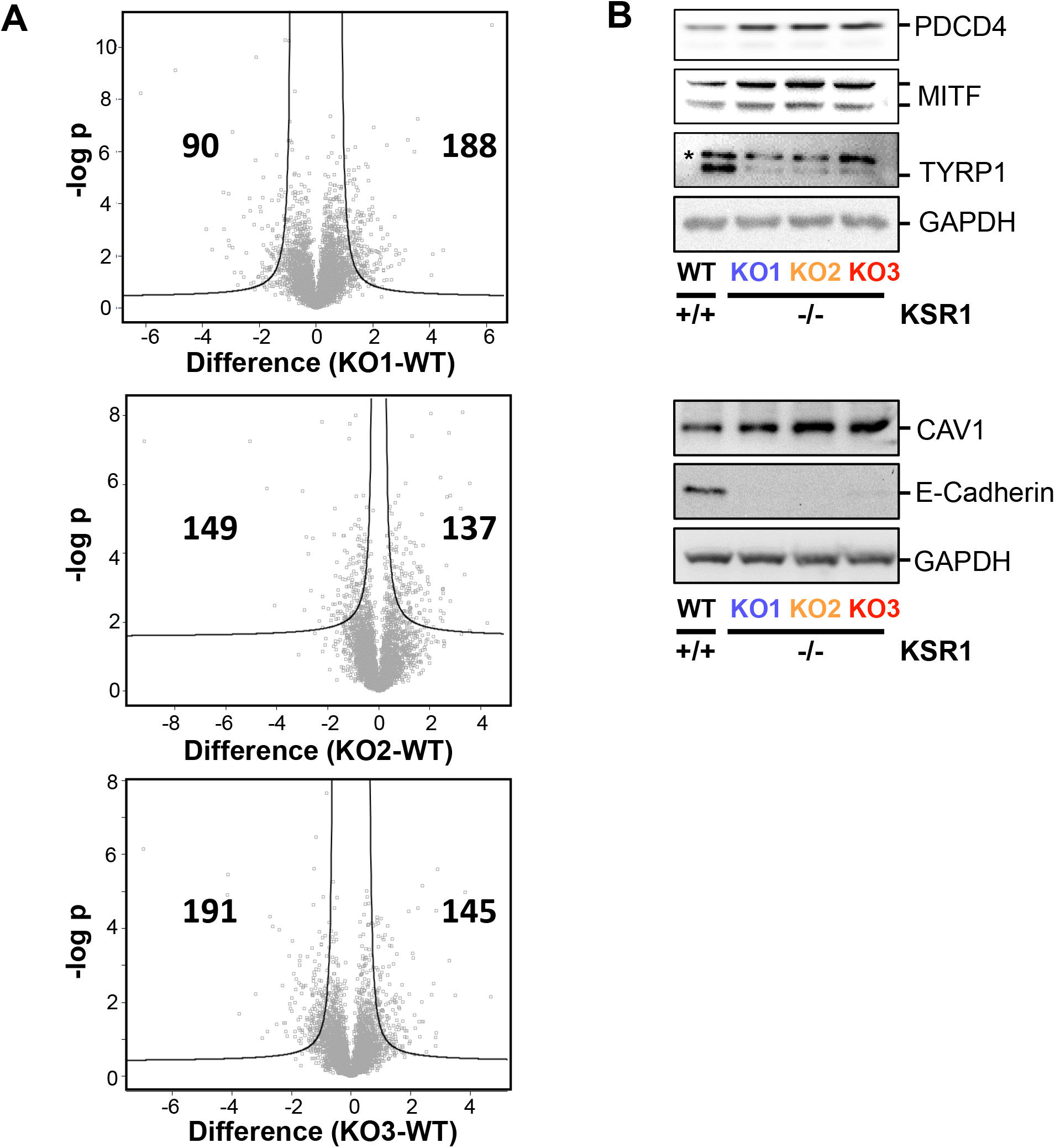
Proteomic expression profiling. **(A)** Volcano plots showing differential protein expression between parental SK-MEL-239 cells and individual KSR1 knockout clones. X-axis, fold difference; Y-axis, -log of p-value. Inset numbers indicate differentially expressed proteins. **(B)** Western blot validation of proteins found differentially expressed by MS based protein expression profiling.

