## Supplemental Table S1 for "Kinase Suppressor of RAS 1 (KSR1) maintains the transformed phenotype of BRAFV600E mutant human melanoma cells"

| Client Sample Name | SK-MEL-239 |
|---|---|
| Sample Code | CL00000490 |
| D8S1179 | 13,14 |
| D21S11 | 29,31.2 |
| D7S820 | 8,10 |
| CSF1PO | 12,12 |
| D3S1358 | 17,17 |
| TH01 | 7,9.3 |
| D13S317 | 12,13 |
| D16S539 | 11,12 |
| D2S1338 | 20,20 |
| D19S433 | 13,13 |
| vWA | 16,17 |
| TPOX | 9,11 |
| D18S51 | 15,18 |
| AMEL | X,X |
| D5S818 | 11,13 |
| FGA | 20,21,22 |
| Database Name | No Hit |

| Client Sample Name | KO1 |
|---|---|
| Sample Code | CL00000491v |
| D8S1179 | 13,14 |
| D21S11 | 29,31.2 |
| D7S820 | 8,10 |
| CSF1PO | 12,12 |
| D3S1358 | 17,17 |
| TH01 | 7,9.3 |
| D13S317 | 12,13 |
| D16S539 | 11,12 |
| D2S1338 | 20,20 |
| D19S433 | 13,13 |
| vWA | 16,17 |
| TPOX | 9,11 |
| D18S51 | 15,18 |
| AMEL | X,X |
| D5S818 | 11,13 |
| FGA | 20,21,22 |
| Database Name | No Hit |

| Client Sample Name | KO2 |
|---|---|
| Sample Code | CL00000493 |
| D8S1179 | 13,14 |
| D21S11 | 29,31.2 |
| D7S820 | 8,10 |
| CSF1PO | 12,12 |
| D3S1358 | 17,17 |
| TH01 | 7,9.3 |
| D13S317 | 12,13 |
| D16S539 | 11,12 |
| D2S1338 | 20,20 |
| D19S433 | 13,13 |
| vWA | 16,17 |
| TPOX | 9,11 |
| D18S51 | 15,18 |
| AMEL | X,X |
| D5S818 | 11,13 |
| FGA | 20,21,22 |
| Database Name | No Hit |

| Client Sample Name | KO3 |
|---|---|
| Sample Code | CL00000494 |
| D8S1179 | 13,14 |
| D21S11 | 29,31.2 |
| D7S820 | 8,10 |
| CSF1PO | 12,12 |
| D3S1358 | 17,17 |
| TH01 | 7,9.3 |
| D13S317 | 12,13 |
| D16S539 | 11,12 |
| D2S1338 | 20,20 |
| D19S433 | 13,13 |
| vWA | 16,17 |
| TPOX | 9,11 |
| D18S51 | 15,18 |
| AMEL | X,X |
| D5S818 | 11,13 |
| FGA | 20,22 |
| Database Name | No Hit |

**Supplementary Table S1. Genotypes of SK-MEL-239 cells and KSR1$^{-/-}$ clones.** Genetic lineage analysis of parental SK-MEL-239 and KSR1$^{-/-}$ single cell clones (KO1-3). Genetic characteristics were determined by PCR-single-locus-technology using the Thermo Fisher, AmpFlSTR® Identifiler® Plus PCR Amplification Kit. In parallel, positive and negative controls were carried out yielding correct results.
